# Bioinspired Stretchable Transducer for Wearable Continuous Monitoring of Respiratory Patterns in Humans and Animals

**DOI:** 10.1101/2022.01.24.477637

**Authors:** Yasin Cotur, Selin Olenik, Tarek Asfour, Michael Bruyns-Haylett, Michael Kasimatis, Ugur Tanriverdi, Laura Gonzalez-Macia, Hong Seok Lee, Andrei S. Kozlov, Firat Güder

## Abstract

We report a bio-inspired continuous wearable respiration sensor modeled after the lateral line system of fish which is used by the fish for detecting mechanical disturbances in the water. Despite the clinical importance of monitoring respiratory activity in humans and animals, continuous measurements of breathing patterns and rates are rarely performed in or outside of clinics. This is largely due to conventional sensors being too inconvenient or expensive for wearable sensing for most individuals and animals. The bio-inspired air-silicone composite transducer is placed on the chest and measures respiratory activity by continuously measuring the force applied to an air channel embedded inside a silicone-based elastomeric material. The force applied on the surface of the transducer during breathing changes the air pressure inside the channel which is measured using a commercial pressure sensor and mixed-signal wireless electronics. We extensively characterized the transducer produced in this work and tested it with humans, dogs, and laboratory rats. The bio-inspired air-silicone composite transducer may enable the early detection of a range of disorders that result in altered patterns of respiration. The technology reported can also be combined with artificial intelligence and cloud computing to algorithmically detect illness in humans and animals remotely, reducing unnecessary visits to clinics.

## 1. Introduction

Respiratory patterns, such as the rate and volume, are known to be important clinical indicators of health both for humans and animals.^1,2^ In human health, monitoring respiration is not only important for diagnosing and tracking respiratory conditions such as sleep apnea, chronic obstructive pulmonary disease, and pneumonia, but also hypoxia, hypercapnia, cardiac arrest, emotional stress, and physical effort.^3–5^ Cretikos et. al. reported that breathing rate in particular is one of the first vital signs to change in response to a problem within the body.^6^ Despite its clinical importance of respiratory rate, respiratory rate is still one of the most poorly monitored vital signs, often relying on infrequent, visual assessments by healthcare professionals (*i.e*. periodically observing the movement of the chest of a patient).^7^ Visual measurements are prone to errors, especially if the patient is aware that their breathing is being monitored and infrequent measurements may miss important events concerning respiratory patterns.^8,9^ Monitoring respiratory activity, such as the breathing rate, is equally, if not more important for animals than for humans; because animals are not able to communicate pain or discomfort.^10^ Research has revealed that variations in respiration (*i.e*., patterns, rate, and depth) in dogs can indicate physical or emotional stress, cardiac, respiratory, and other health-related problems(including heat exhaustion).^11^ In some animals, such as dogs, measuring breathing activity manually by a human or in an unfamiliar environment may also result in unreliable measurements due to emotional factors, leading to poor clinical decisions and outcomes.^12^ Monitoring breathing is also equally important in laboratory animals (such as rats) during various procedures including anesthesia but the existing instruments are expensive and not precise.^13^

To address the problems concerning manual, infrequent, or unreliable assessments, wearable continuous breathing monitors have been reported, although the primary focus to date has been human health.^14–17^ Limitations related to performance longevity and discomfort have prevented the wide-scale adoption of continuous breathing monitors by the health systems or consumers, however.^18–22^ Wearable systems for continuous respiratory monitoring can be primarily divided into two categories: sensors that measure breathing by sensing *i)* the airflow through the nose and mouth^23–27^ and; *ii)* the movement/volumetric expansion and contraction of the chest/abdomen of the subject wearing the instrument.^28–33^

Wearable continuous measurement of airflow due to breathing has been limited to human health and is commonly achieved by exploiting the difference between the temperature and moisture content of the exhaled breath and the external environment, which can be captured using temperature or humidity sensors.^34,35^ Although low-cost and simple to use, the sensitivity of temperature and moisture sensors decreases when the difference in temperature or relative humidity between the external environment and exhaled breath is small. When sufficient cooling is not provided, temperature sensors suffer from the accumulation of heat, resulting in loss of function and drift.^36^ Similarly, for humidity sensors, sufficient ventilation, and protective layers are required (increasing complexity and cost) to prevent accumulation of moisture and condensation which reduce function and cause drift over time.^37^ Furthermore, because direct monitoring of flow requires sensors to be worn on the face, they cannot be worn discreetly.

Measuring breathing using instruments attached to the chest allows continuous monitoring of respiratory activity in both humans and animals without directly monitoring the airflow. Accelerometers have been used to measure breathing patterns both in humans and animals by way of capturing the movement of the chest during cycles of inhalation and exhalation.^38,39^ Accelerometers have been used to capture these movements; however, they are susceptible to motion artifacts and do not typically produce large, detectable signals – *i.e*., the signal-to-noise ratio is poor. Various groups, including our laboratory, have also demonstrated the use of wearable strain sensors to continuously measure the volumetric expansion of the chest/abdomen due to breathing.^40–45^ Notable works include strain sensors based on textiles, conductive polymer composites, or microfabricated adhesive patches with metal electrodes.^46–48^ These sensors require either complex manufacturing technologies or rely on custom formulations of materials that cannot be purchased off the shelf. These two critical issues lead to poor reliability and increased costs, eventually slowing clinical translation and reducing the applicability of the devices produced in real-world settings.

In this work, we present a low-cost, wireless, and stretchable wearable transducer for continuous monitoring of respiratory activity in humans and animals (in this study dogs and rats). The <u>a</u>ir-<u>si</u>licone composite transducer (ASiT), inspired by the lateral line system of fish,^49^ consists of silicone rubber with an embedded millimeter-scale air channel (**Figure 1**). The air pressure inside the channel changes when a force is applied across the surface of the device which is monitored with a commercially available solid-state air-pressure sensor. ASiT is strapped to the chest/abdomen of the subject (like a rubber band) and responds to volumetric expansion and contraction during cycles of inspiration and expiration. We have optimized the ASiT in laboratory experiments to produce a reliable system and characterized its performance in a series of human and animal (*i.e*., dog and rat) experiments for continuous monitoring of respiratory activity.

**Figure 1.**
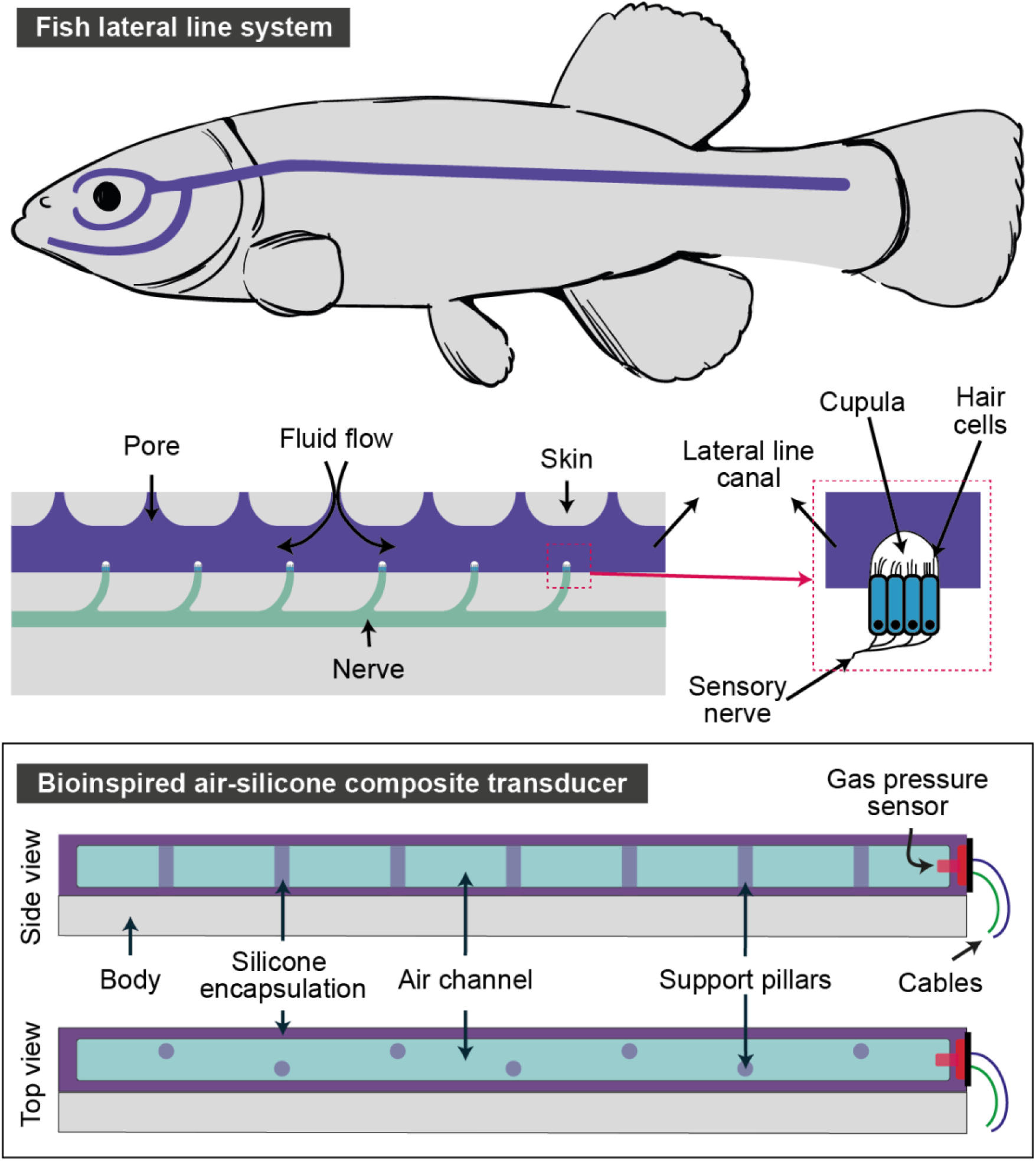
Schematic representation of the fish lateral line system and its cross-sectional structure consisting of a fluid canal, sensory nerves, and hair cells to detect the pressure changes in water. Schematics illustration of the bioinspired air-silicone composite transducer (ASiT) which contains an air channel encapsulated with a thin layer of silicone elastomer with support pillars also made of the same material. The pressure applied on the silicone air channel changes the air pressure within the channel, which can be measured using a differential gas pressure sensor and digitized using electronics.

## 2. Results and Discussion

### 2.1. Fabrication of the bioinspired air-silicone composite transducer

The lateral line system of fish contains mechanoreceptors called hair cells that are sensitive to mechanical stimuli, such as water pressure gradients.^50^ In the lateral line system, groups of hair cells are encapsulated with a long and thin canal extending along the trunk and can sense tiny changes in pressure in the water (Figure 1). ASiT was developed taking inspiration from the lateral line system of fish for sensing external pressures. The silicone-encapsulated air channel mimics the lateral line canal, the gas pressure sensor mimics the hair cells, and cables mimic the sensory nerves

We fabricated the bioinspired air-silicone composite transducer in six steps (supporting information **Figure S1**): (i) We first produced the top silicone layer of the transducer using a 3D-printed mold made of polylactic acid (PLA). (ii) After partially curing the top silicone layer for two hours at room temperature, we prepared the bottom silicone layer by pouring liquid (uncured) silicone into a rectangular 3D-printed mold, 1 mm in height. (iii) We left the partially cured top and uncured bottom silicone layers to cure together for another two hours at room temperature to form a monolithic part with an internal air channel. (iv) A small silicone-based hollow tube with an outer diameter of 3mm was connected to the air channel; a blunt syringe needle was used to punch a hole through the walls of the monolithic silicone structure and guide the tube into the air cavity. We used Sil-Poxy™ silicone adhesive (by Smooth-On, Inc.) to seal the edges of the tube to prevent air leakage from the air cavity. (v) The other end of the silicone tube was connected to a commercially available gas pressure sensor (vi) Finally, we attached the ASiT to a silicone-based wearable stretchable harness containing a wireless electronics unit to form the entire device. The electronics unit was custom designed to acquire the respiration signal and to transmit to a nearby device over Wi-Fi. The sticky nature of silicone improved adhesion especially on the clothing and animal fur, preventing the harness and transducer from slipping on the subjects (**Figure 2**).

**Figure 2.**
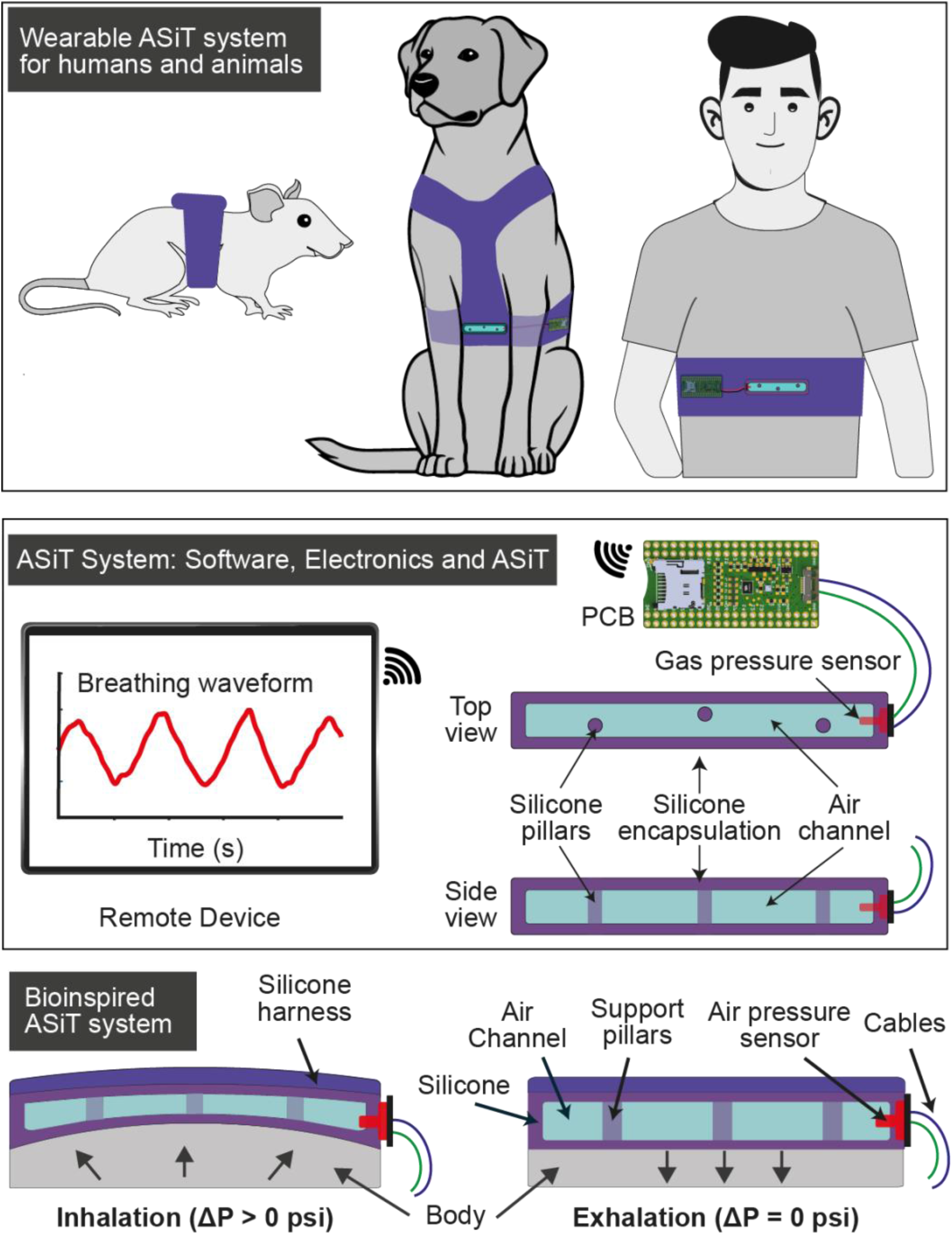
Schematic presentation of the wireless wearable ASiT system for monitoring respiratory activity. Compression of ASiT during inhalation increases the air pressure inside the silicone air channel; exhalation brings the air pressure back to initial pressure creating a self-referenced respiration signal. The changes in the air pressure inside the air channel are captured with a commercially available gas pressure sensor, allowing continuous and real-time monitoring of respiratory activity in humans and animals. (PCB: Printed Circuit Board)

### 2.2. Characterization of the performance

The ASiT senses breathing by measuring the air pressure within the air channel inside the stretchable silicone structure which deforms during breathing due to the force applied to its outer surface by the expanding chest. To understand the impact of geometry to create a transducer with high sensitivity (*i.e*., large pressure differential under applied force), we designed five ASiTs with varying geometries with each having three additional geometrical variants (see supporting information **Figure S2** and **S3** for further details). A sensitive transducer enables the detection of respiration patterns of low intensity (*e.g*., panting in dogs), hence it is an important feature for respiration monitoring. With Designs (1-5) and variants (a-c), we primarily tested the role of shape, air volume, and thickness of the silicone walls of the stretchable structure on sensitivity (**Figure 3**). To simulate the forces exerted during repeated cycles of inhalation and exhalation, we used a programmable force testing system (MultiTest 5-xt by Mecmesin UK, Figure 3A) for increased precision. We applied a periodically varying force signal between 0 N and 5 N to the transducers and recorded the change in air pressure within the air channel inside the silicone structure (see Figure 3B and C for representative air pressure and force waveforms obtained from Design 5 variant ‘a’; see supporting information **Figures S4** and **S5** for pressure and force waveforms for each design and variant). Figure 3D compares the performance of each design and variant under an applied force of 5 N; a larger pressure differential produced under the same force indicates better sensitivity. All geometries produced a directly proportional relationship between the force applied and the pressure measured. We have discovered that two parameters play an important role in defining sensitivity: the thickness of the walls of the stretchable silicon structure containing the air channel and the volume of the air channel. Thicker walls reduce the ability of the silicone structure to deform under applied force, resulting in lower sensitivity. Lowering the volume of the air cavity increases sensitivity because of the increased compression for the same force.

**Figure 3.**
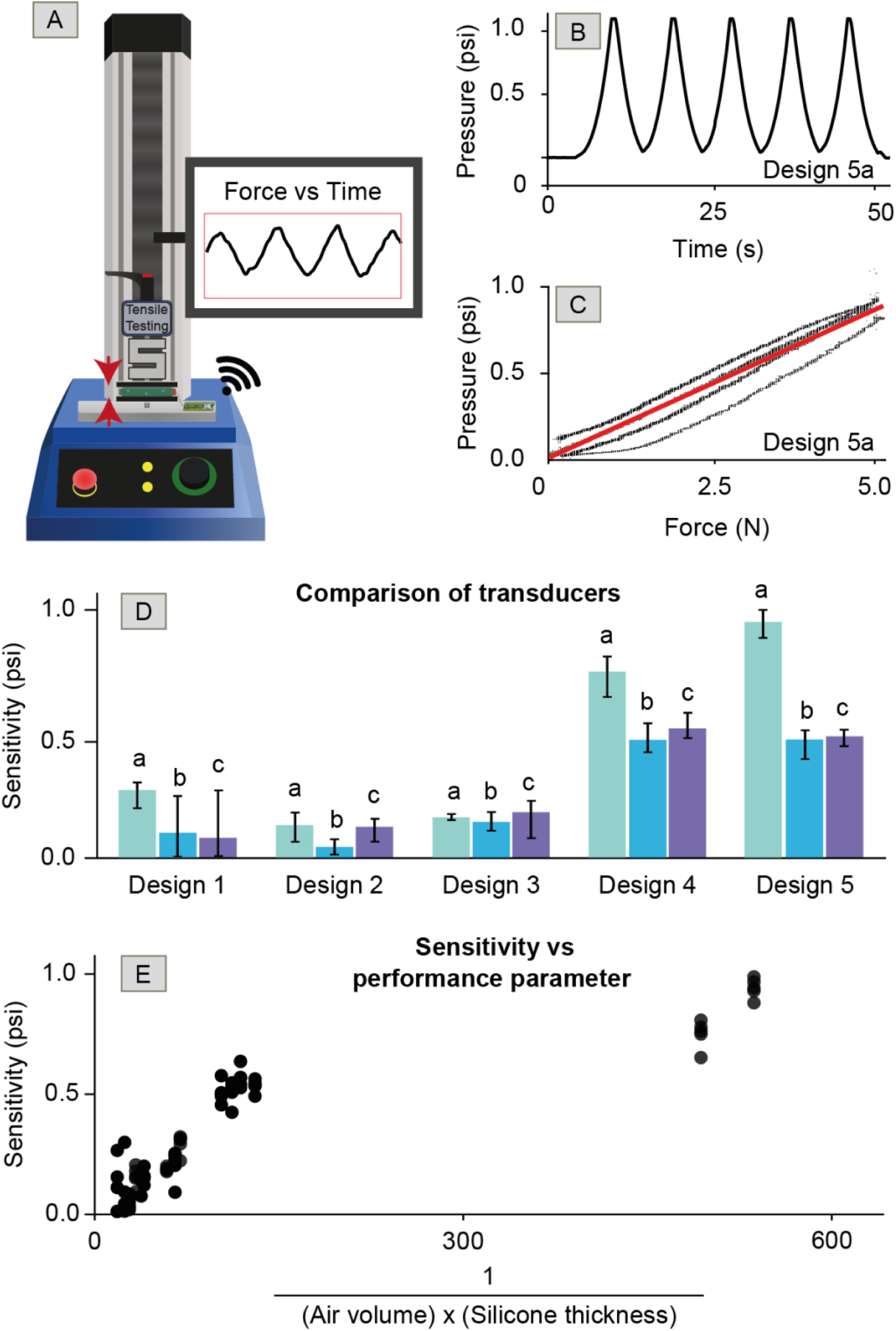
Comparison of the performance of transducer design variants. Transducer *Design 1* has a large-volume air channel, *Design 2* is equivalent in size to *Design 1* but with a lower volume air channel and thicker (silicone) walls, *Design 3* is equivalent in size to *Design 2* but uses three connected air channels with the same dimensions as the channel in *Design 2, Design 4* is identical to *Design 2* but with a thinner wall thickness. *Design 5* is identical to *Design 4* but includes small columns for structural support. **A)** Schematic illustration of the setup used for mechanical characterization of the transducers. A cyclic force of 0-5N was applied on the transducers mimicking breathing and the air pressure inside the air channels was measured. **B)** A representative air pressure vs time graph was obtained from *Design 5a* when a cyclic force of 0-5N was applied for five cycles. **C)** A representative air pressure vs force waveform obtained from *Design 5a*. **D)** Sensitivity (*i.e*., pressure differential) of each Design and their three variants with small alterations when a force of 5 N was applied (n=5). **E)** Plot showing the correlation between sensitivity and the geometrical metric derived by us that can be used for guiding design decisions for improved sensitivity. Error bars: Standard deviation. Descriptions for designs can be found in supporting information Figure S3.

Because of these factors, Designs 4a and 5a produced the highest sensitivity among all geometries tested with Design 5a producing the higher overall sensitivity. Although Design 4a and 5a were nearly identical in terms of the wall thickness and volume of the air cavity, Design 5a produced a higher sensitivity in comparison to 4a. This is because Design 4a did not provide enough mechanical strength to prevent the air channel from collapsing on itself which was improved in Design 5a by placing small support pillars inside the channel. Using these results, we created a performance parameter (*PP*) that defines the relative sensitivity of an ASiT:

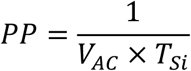

where V_AC_ is the volume of the air channel and *T*_*Si*_ is the thickness of the walls of the stretchable silicone structure. Plotting *PP* against sensitivity also confirms the positive correlation between these two metrics (Figure 3E).

### 2.3. Monitoring breathing of humans

We slightly modified Design 5a to produce a wearable form factor for respiration monitoring (see supporting information **Section S1**) and integrated the ASiT onto a silicone harness, creating a monolithically integrated stretchable wearable device (**Figure 4A**). During inhalation, the expanding chest presses the ASiT against the harness increasing the pressure inside the air channel. The air pressure drops back to the initial levels when the subject exhales. By using a less stretchable silicone formulation (Dragon Skin Ⓡ20 instead of Ecoflex™ 00-30), the stretchability of the harness can be reduced to achieve larger compression against the ASiT hence a larger pressure differential within the air channel during breathing.^51^

**Figure 4.**
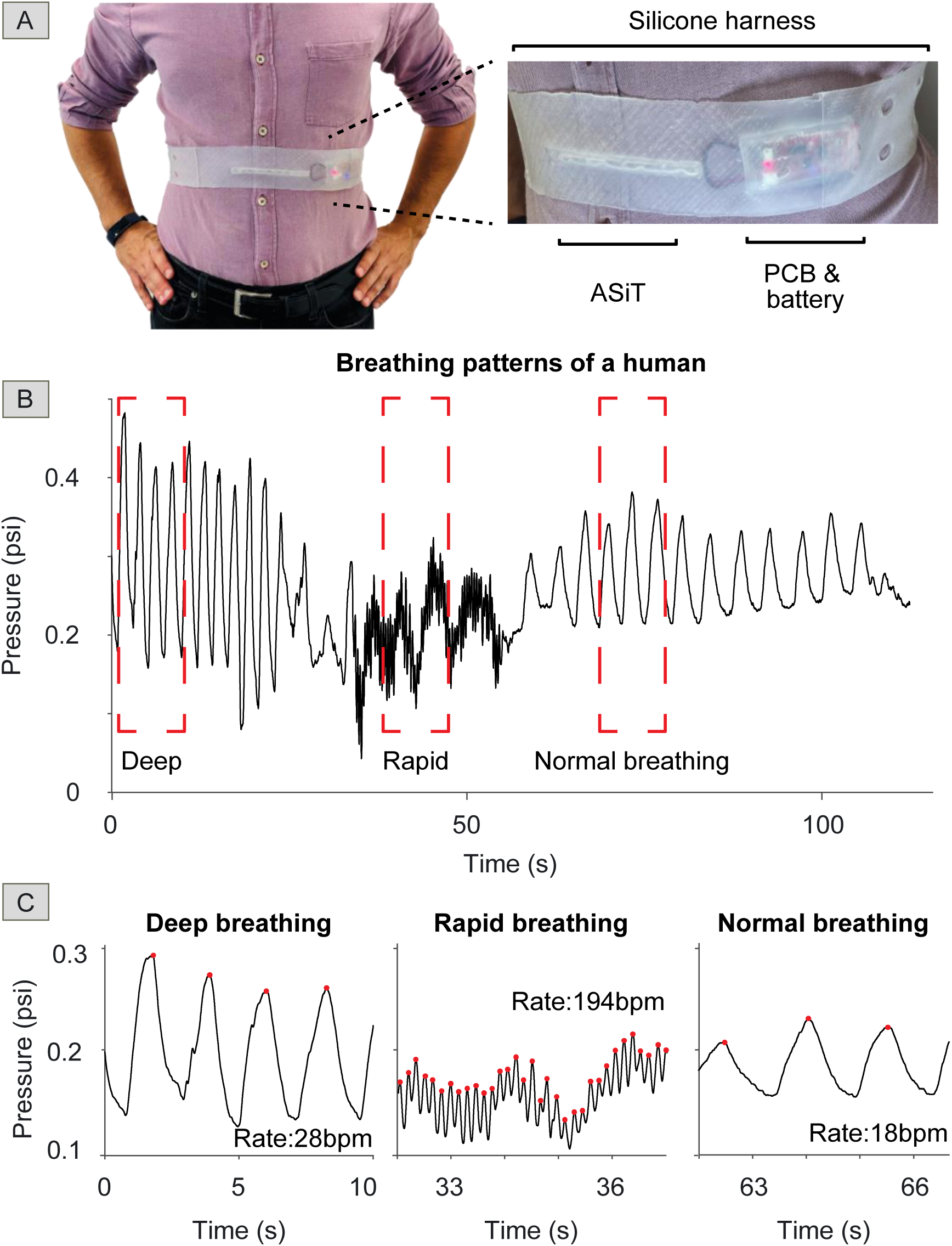
Human experiments. **A)** Photographs of the wearable ASiT system worn by a healthy volunteer. **B)** The respiratory activity of the volunteer was captured using the ASiT in a controlled experiment. **C)** Waveforms showing different patterns of breathing. Breathing rates (breaths per minute – bpm) were estimated automatically using a custom algorithm developed in this work.

Using the wearable ASiT system, we were able to monitor the respiration of a healthy human volunteer continuously (**Figure 4B**). We were able to capture deep, rapid, and normal breathing patterns using the wearable ASiT system with high clarity. We also developed an algorithm for automatic real-time detection of breathing rates to eliminate manual counting of peaks (supporting information **Section S2 and S3**). The algorithm can detect the locations of peaks on the waveform captured where inhalation (high peak) and exhalation (low peak) occur and count the number of peaks every 15 seconds. We have applied this algorithm to the waveform shown in Figure 4B and the results are shown in **Figure 4C**. Each high peak is marked with a red point automatically by our peak detection algorithm. The performance of the algorithm developed was also evaluated by manually counting the breathing rates and comparing to the rates estimated by the algorithm. The algorithm predicted the correct respiration rates for the human subject during normal (18 <u>b</u>reaths <u>p</u>er <u>m</u>inute - bpm), deep (28 bpm), and rapid breathing (194 bpm).

### 2.4. Monitoring breathing of dogs

To demonstrate the versatility of the wearable ASiT system across different animal species for monitoring respiratory activity, we modified the harness designed for humans for use with dogs (**Figure 5**). We produced a silicone-based flexible and stretchable harness (modeled after conventional dog harnesses commonly used by dogs) with a monolithically integrated ASiT for increased reliability and fitting. The ASiT was located on the section of the harness in contact with the chest of the dog. Our harness design (Figure 5A) prevented the harness from moving along the torso of the dog during activity, ensuring consistent measurements with minimal motion artifacts. We tested the wearable ASiT system with a one-year-old Labrador Retriever detection dog that was trained for detecting explosives. We chose to work with an explosive detection dog instead of a pet because they are highly trained to follow instructions (sitting, standing, *etc*.), important for testing the ASiT system in a controlled fashion.

**Figure 5.**
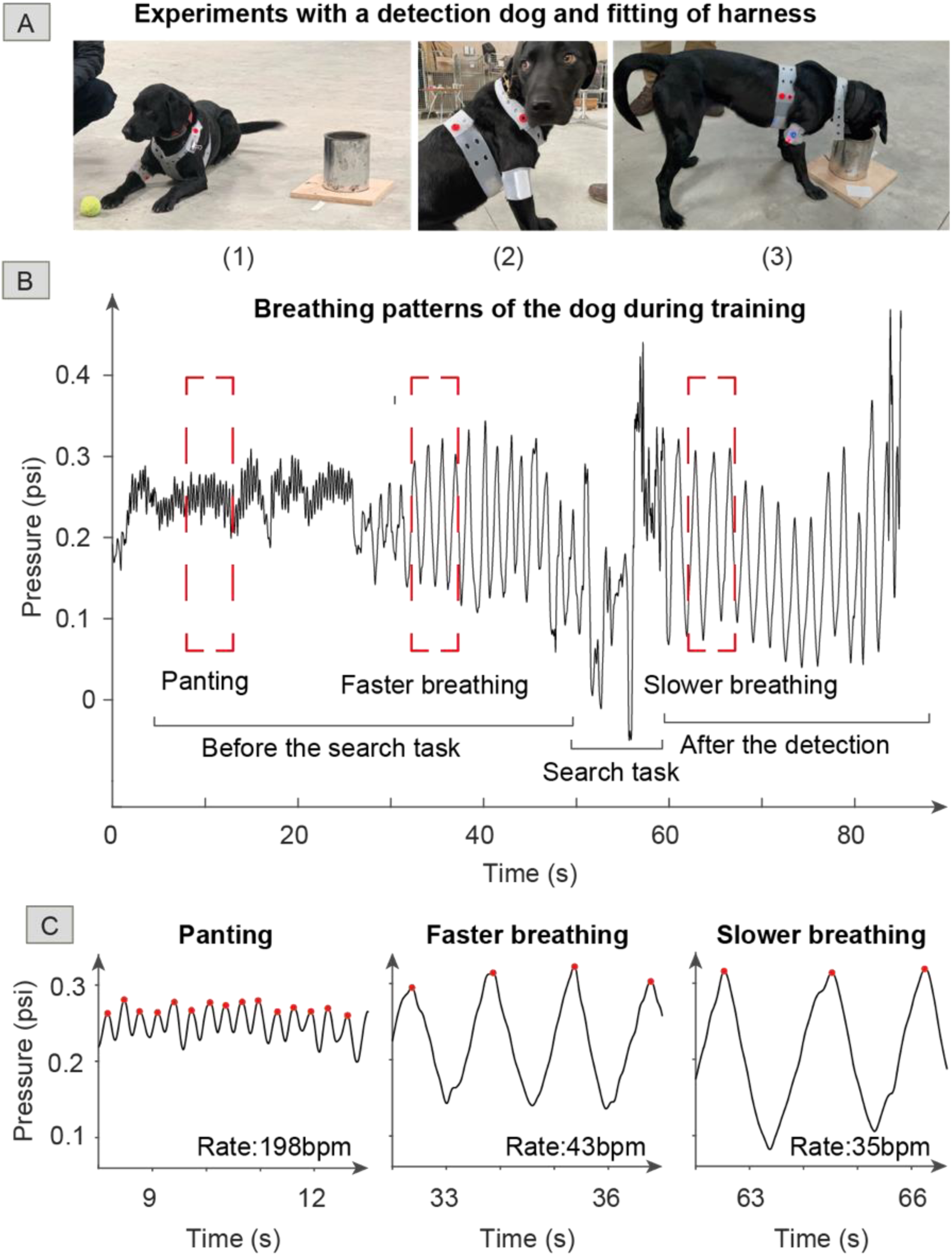
Dog experiments with a Labrador Retriever. Recording of breathing data of a detection dog during training. **A)** Photographs of the wearable ASiT system worn by a detection dog while it was (1) resting after a search task, (2) sitting before a search task, (3) indicating a target odor by freezing in anticipation of a reward. **B)** Respiratory waveform acquired from the detection dog during a training task; the recording shows panting and normal breathing before and after the detection event. **C)** The patterns of breathing while panting, before the search task, and after the detection event of the dog.

We captured the respiratory activity of the subject during a training session which included rest, search, and play (**Video V1**). The breathing waveform recorded is shown in Figure 5B. The ASiT system could successfully capture breathing and panting (high-frequency breathing, generally with the mouth open, that helps with cooling) when the dog was stationary. The automated detection algorithm developed previously could also detect the peaks in the respiration waveform and measure the rate of breathing and panting (Figure 5C). The wearable ASiT system, however, was unable to detect the patterns of breathing when the dog was in fast motion. In this training exercise, the dog first panted for approximately 25 seconds with a rate of 198 bpm and switched to regular breathing (43 bpm) until the search task was initiated by the handler (at ∼50-second mark). After a short journey of 10 meters lasting five seconds, the detection dog identified a target scent that was placed inside a metal canister and froze as an indication behavior for about 30 seconds (until rewarded with a toy). Upon detection of the target scent, the respiration rate dropped from 43 bpm to 35 bpm, a 20% decrease while the depth of breathing increased 15%. Although it is not exactly clear why the respiration rate has dropped after detecting a target scent, we speculate this may be related to the expectation of a reward (in the form of a toy).^52–55^ More research is needed to understand this phenomenon with a larger sample size (in this study, n=1).

After the training tasks were completed, the dog was taken away to a quiet area for resting. This allowed us to continue monitoring the respiratory activity of the dog during rest (**Figure 6**). We noted shifts in the baseline in the signal recorded because of the change in the baseline force exerted on the transducer in different positions (sitting, lying down, standing) during rest (Figure 6A). These shifts in the baseline, however, did not visibly impact the quality of the recording and can easily be removed digitally.^4^ The recording of the respiratory activity at rest revealed several interesting patterns: we observed that during a period lasting a few minutes (Figure 6B), a full cycle of breathing included two identifiable peaks (one large and one small) highlighted in the red in Figure 6B. Approximately 20 minutes later, we also observed another pattern. Here, a deep breath was followed by a 10-second pause after which the dog went back to breathing with a lower frequency, but higher depth compared to before the pause (Figure 6C). Such patterns of breathing are likely not related to any pathologies hence normal. But the capability of detecting small variations in breathing may enable the identification of biomarkers for various diseases or the emotional state of the animal which can be detected algorithmically using the software.

**Figure 6.**
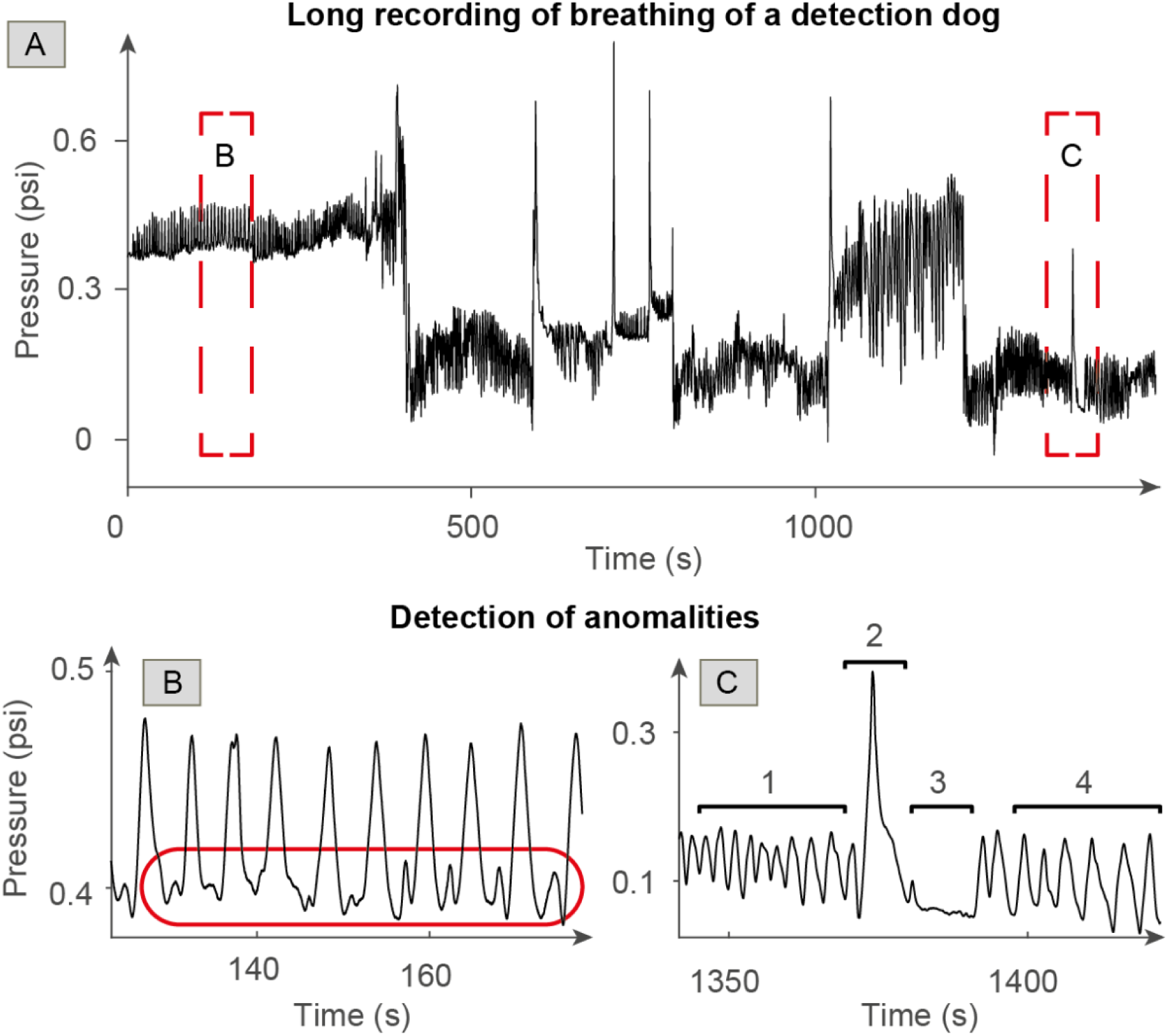
**A)** The recording of breathing of the detection dog during prolonged resting. Shifts in the baseline are due to a change in posture which applies a different baseline force on the transducer. **B)** Unusual respiratory patterns captured from the dog (marked with the red circle) while resting in a quiet area. **C)** ASiT captures a breathing pattern consisting of fast (1), deep (2), paused (3), and slow breathing (4). Breathing rate decreased after the deep-paused breath.

### 2.5. Monitoring breathing of rats

Because monitoring the respiratory activity of laboratory animals is essential during various lab procedures and to evaluate the performance of ASiT in small-sized animals, we also conducted experiments with a Sprague Dawley laboratory rat (**Figure 7**). We redesigned the silicone harness, integrated with an ASiT and electronics, to fit on the body of the rat (Figure 7A). Since the chest of the rat was smaller than that of humans and dogs, we also had to use a smaller air-silicone transducer (Design 5a with the dimensions of 40 × 60 × 5 mm^3^).

**Figure 7.**
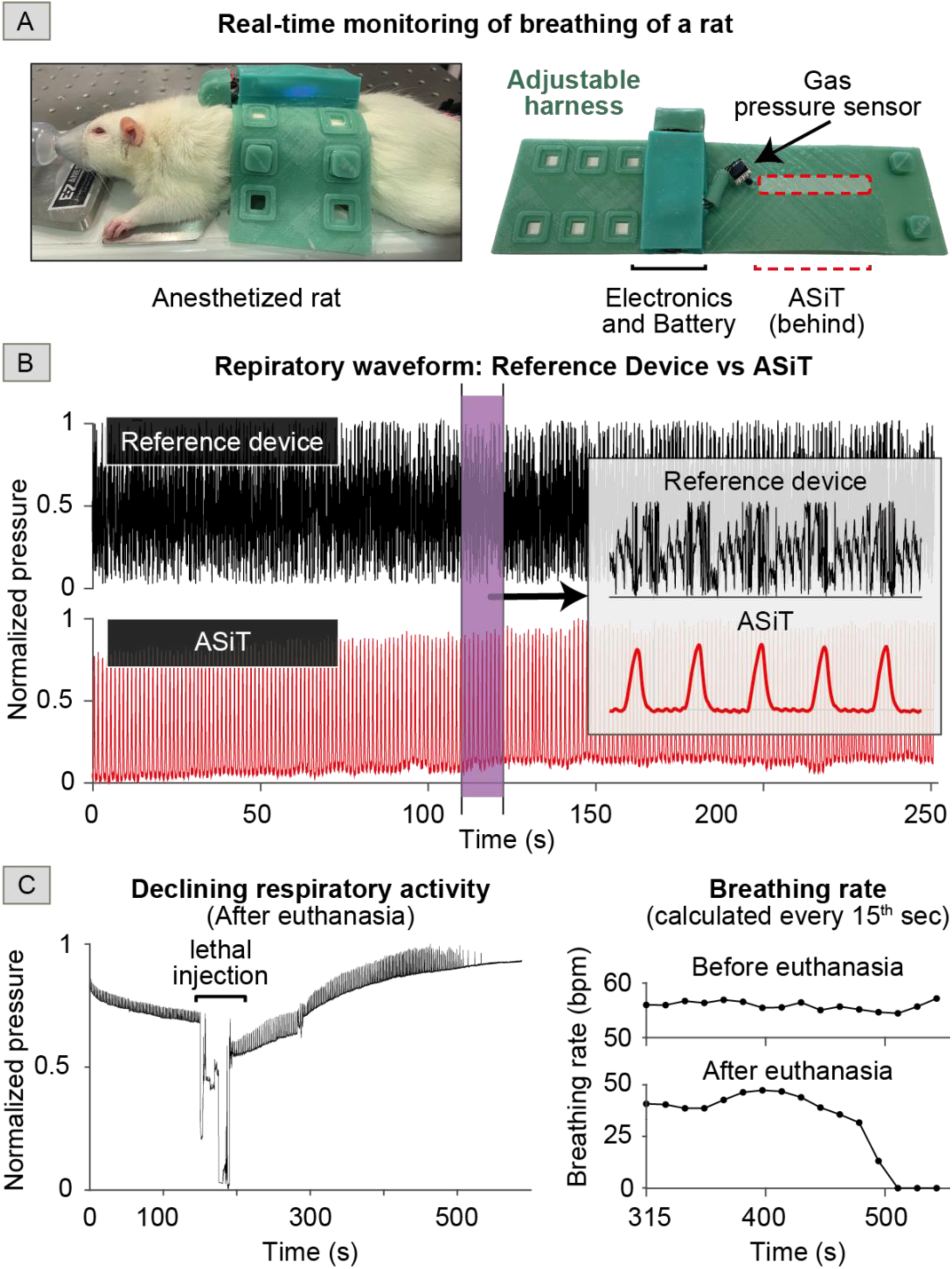
Rat experiments with a Sprague Dawley laboratory rat under anesthesia. **A)** Photographs of the wearable ASiT system worn by a laboratory rat and the isometric view of the harness and its elements. **B)** Respiratory waveforms were captured using a commercial (reference) laboratory instrument and ASiT at the same time. ASiT produced substantially clearer signals in comparison to the reference device. ASiT also appear to pick up the cardiac activity of the rat indicated by the small spikes in the plot shown in the inset. **C)** Respiratory activity during the pre-planned euthanization of the rat with a lethal dose of intraperitoneal while anesthetized. Increasing baseline pressure indicates loss of muscle tone. A sharp decrease in breathing rate was also observed a few minutes after the injection.

We recorded the respiratory activities of the rat under anesthesia using ASiT (**Video V2**). Simultaneously, we also used a commercial device (Small Animal Physiological Monitoring System by Harvard Apparatus) for measuring breathing that is used routinely in animal research laboratories worldwide. In terms of the quality of the signals acquired, ASiT performed substantially better in comparison to the commercial device which produced a noisy signal (Figure 7B). The signal acquired from ASiT is sensitive and noise-free to the degree that we were also able to pick up the activity of the rat’s heart as indicated by the small spikes shown within the breathing patterns in the inset of Figure 7B. We also recorded the breathing activity of the rat during a planned euthanization (Figure 7C). Whilst anesthetized with isoflurane, the rat was euthanized with a lethal intraperitoneal injection of sodium pentobarbital. After the injection, the baseline pressure within the ASiT slowly increased as the rat lost muscle tone causing it to lean more on the sensor. We also calculated the breathing rate of the rat before and after the lethal injection. Approximately three minutes after the injection, breathing stopped. The ASiT was able to accurately record changes in the breathing rate, as both the rate and amplitude gradually decreased to zero. These results demonstrate that the wearable ASiT system can be miniaturized for use with small animals.

## 3. Conclusions

The air-silicone composite transducer produced in this work was fabricated with commonly available prototyping tools (such as 3D printing) and materials (such as silicone rubbers), easily accessible to most academic and industrial laboratories in the world. The materials used were relatively low-cost, costing ∼$40 for a single laboratory prototype ($35 for electronics, $5 for 3D printed molds, and silicone rubbers for each). The costs could be further reduced by using an alternative gas pressure sensor, which was the most expensive component used in our design ($25 / sensor), or ordering components in larger quantities.

The method presented to design and produce the transducers is versatile and can be easily modified to provide excellent fitting regardless of the size or the species of the subject. ASiT, when used as a wearable sensor for continuous monitoring of respiratory activity, produces an information-rich, time-series waveform that could be algorithmically processed to extract physiological data (see supporting information **Section S3**). ASiT does not require calibration and is sufficiently sensitive to detect all forms of respiratory activity such as shallow breathing or panting.

The ASiT has at least three limitations: i) Although wearable ASiT system is soft, flexible, and non-invasive, it requires direct and continuous contact with the body, which may cause discomfort for some users when used for a long time. ii) The performance of the transducer drops due to the deformation of the silicone-based air channel when the subject is in motion. Because of this, the highest quality respiratory data are collected when the subject is stationary. iii) The custom-made wireless electronics board used in this study has high power requirements (up to 300mA especially during Wi-Fi transmission) and the 500 mAh LiPo battery used in our experiments provides a runtime of max. 2.5 hours. Although this duration was sufficient for our experiments, for clinical applications, this may be too short. Hence a complete redesign of the electronics board is likely needed for real-world applications.

Using the wearable ASiT system, the digital respiratory data collected from animals and humans can be transmitted to a nearby mobile device for algorithmic processing and cloud storage. The data on the cloud can be shared with health systems, such as medical or veterinary practices, on-demand when anomalies are detected automatically. The digital and cloud capabilities of the wearable ASiT system are also compatible with emerging technologies, such as blockchain-based smart contracts that can completely automate care.^56^ Furthermore, monitoring companion animals in their usual settings would improve the quality of the physiological data collected for a more precise assessment of health and wellbeing. The wearable ASiT system, when combined with other technologies for physiological monitoring,^5^ would allow early diagnosis of a wide range of animal and human diseases, reducing unnecessary visits to clinics while improving outcomes.

## 4. Methods and Materials

### Human and animal experiments

Based on the guidelines at Imperial College London, the project team performed a detailed risk assessment to identify the hazard and risks associated with non-invasive human and dog experiments. All experiments were conducted on healthy subjects. The dog experiments were performed in collaboration with RFA Security Ltd with their detection dogs. The dog that participated in the project was closely observed for any signs of stress by the owner and project team. One female Sprague Dawley (CD) rat (330g) was anesthetized using isoflurane (inducted in a chamber and then moved to a facemask), and the customized harness was placed around the thorax. Readings were then taken at various concentrations of isoflurane (2-0.5%). The procedure on the rat was performed in accordance with the 1986 Animal (Scientific Procedures) Act, under approval from the UK Home Office.

### Transducer and harness design

We measured the air pressure exerted in the ASiT around 7kPa (1psi) from an applied 5N force using a Mecmesin Universal testing machine MultiTest 5-xt (Figure 3A). The intrapleural pressure of a human can increase to 8kPa in deep breathing for humans.^57^ To measure the change of the air pressure inside the transducer, we used a commercially available, differential gas pressure sensor (HSCDLNN001PDAA3) from Honeywell Sensing and Productivity Solutions. The sensor has high stability and reliability and can detect gas pressures from −1 psi to 1 psi (around 7kPa).

The silicone harnesses were designed to conform to the body of the subject. The harnesses designed for rats and dogs had three pairs of holes for adjusting tightness. ASiT was placed on the side of the harness contacting the body of the subject. The harness for dogs included an additional belt-like silicone extension to secure the harness around the neck of the animal to improve fitting and reduce slipping of the harness along the body of the animal when in motion.

### Electronics & software

In our experiments, we used an ESP32 microcontroller-based electronics module by SparkFun with a built-in voltage regulator and battery charger. This module worked with a lithium-ion battery (3.7V, 500mA) lasting approximately 2.5 hours with a Wi-Fi transmission on. ESP32 was chosen as the preferred microcontroller because of its fast Xtensa dual-core 32-bit LX6 compute units and built-in wireless communication features such as Bluetooth and Wi-Fi which accelerated the time for implementation. To complement the capabilities of the purchased Sparkfun ESP32 module, we designed an additional printed circuit board (PCB) to add a micro-SD card reader for data storage (see supporting information **Figure S6**). Another PCB (Sensor-PCB) was also designed and produced to include additional electronics to acquire more precise measurements from the gas pressure sensor (see supporting information **Figure S7**). The Sensor-PCB contained a 16-bit analog-to-digital converter (ADS8326 by Texas Instruments) to improve the sensitivity of the recordings from the pressure sensor. The readings from the ADS8326 were further improved by using a dedicated voltage reference (REF2033 by Texas Instruments). A flexible flat cable (0150180105, CABLE FFC 10POS 0.50MM 6”) was used to connect the ADS8326 chip to the ESP32 module. The pressure readings generated by the gas pressure sensor were stored on the SD and at the same time, transmitted to a nearby PC over Wi-Fi.

The gas pressure sensor was sampled at 1kHz and smoothened using a moving average filter with a smoothing constant of 50 samples to construct the respiratory signals. We designed a user interface on MATLAB (see supporting information **Figure S8**) for remote operation and data communication with the electronics unit. MATLAB was also used for data processing and analysis. All electronic components were purchased from Digikey, PCBs and stencils were sourced from AISLER and Elecrow. Schematics and PCBs were designed in EAGLE. The ESP32 was programmed in PlatformIO embedded in Visual Studio Code using Arduino libraries.

### Fabrication of molds for the production of silicone components

All molds used in the experiments for producing silicone-based components were designed in Dassault Systèmes SOLIDWORKS Corp. Polylactic Acid (PLA) filaments were purchased from 3dgbire and RS Components and used as the thermoplastic for 3D printing. The molds were 3D printed using Raise3D N2 Plus and BCN3D Sigmax R19 printers. We used the silicone elastomer Ecoflex™ 00-30 (produced by Smooth-On, USA and ordered from Bentley Advanced Materials, UK) in the fabrication of the ASiT by mixing Part A and Part B with a ratio of 1:1 by weight. Once mixed, the silicone mixture was degassed for 15 min in a vacuum chamber to remove any air bubbles before pouring molding at room temperature. The harnesses were produced using the same silicone-based fabrication method as ASiTs, but a less stretchable elastomer formulation (Dragon Skin Ⓡ20 by Smooth-On) was used instead.

### Algorithmic detection of breathing rate

We have developed a custom breathing rate detection algorithm on MATLAB for the analysis of the breathing waveforms. The details of the algorithm used can be found in the supporting information **Section S2**.

## 5. Acknowledgements

Firat Güder and Ugur Tanriverdi would like to thank Imperial College Department of Bioengineering. Firat Güder and Yasin Cotur would like to thank the Institute for Security Science and Technology (funding under Champions Fund) and financial support. Yasin Cotur would like to thank the Turkish Ministry of Education and EPSRC IAA. Selin Olenik acknowledges the Imperial President’s Ph.D. Scholarship. Michael Kasimatis acknowledges EPSRC DTP (Reference: 1846144). Firat Güder and Tarek Asfour acknowledge Imperial College Centre for Processable Electronics and Innovate UK (grant reference: 10004425). Firat Güder and Laura Gonzalez-Macia would like to thank the Bill and Melinda Gates Foundation (Grand Challenges Explorations scheme under grant number: OPP1212574) and the US Army (U.S. Army Foreign Technology (and Science) Assessment Support (FTAS) program under grant number: W911QY-20-R-0022) for their generous support. Work in the Kozlov lab is funded by the Wellcome Trust (214234/Z/18/Z), The US Army Research Laboratories, and the Centre for Blast Injury Studies at Imperial College London. We would like to thank Merve Cirisoglu Cotur for her help with the illustrations. We would also like to thank Tony Foster and Peter Watson from RFA Security for their help with the animal experiments.

