## Supplementary information for "Bioinspired Stretchable Transducer for Wearable Continuous Monitoring of Respiratory Patterns in Humans and Animals"

### Figures

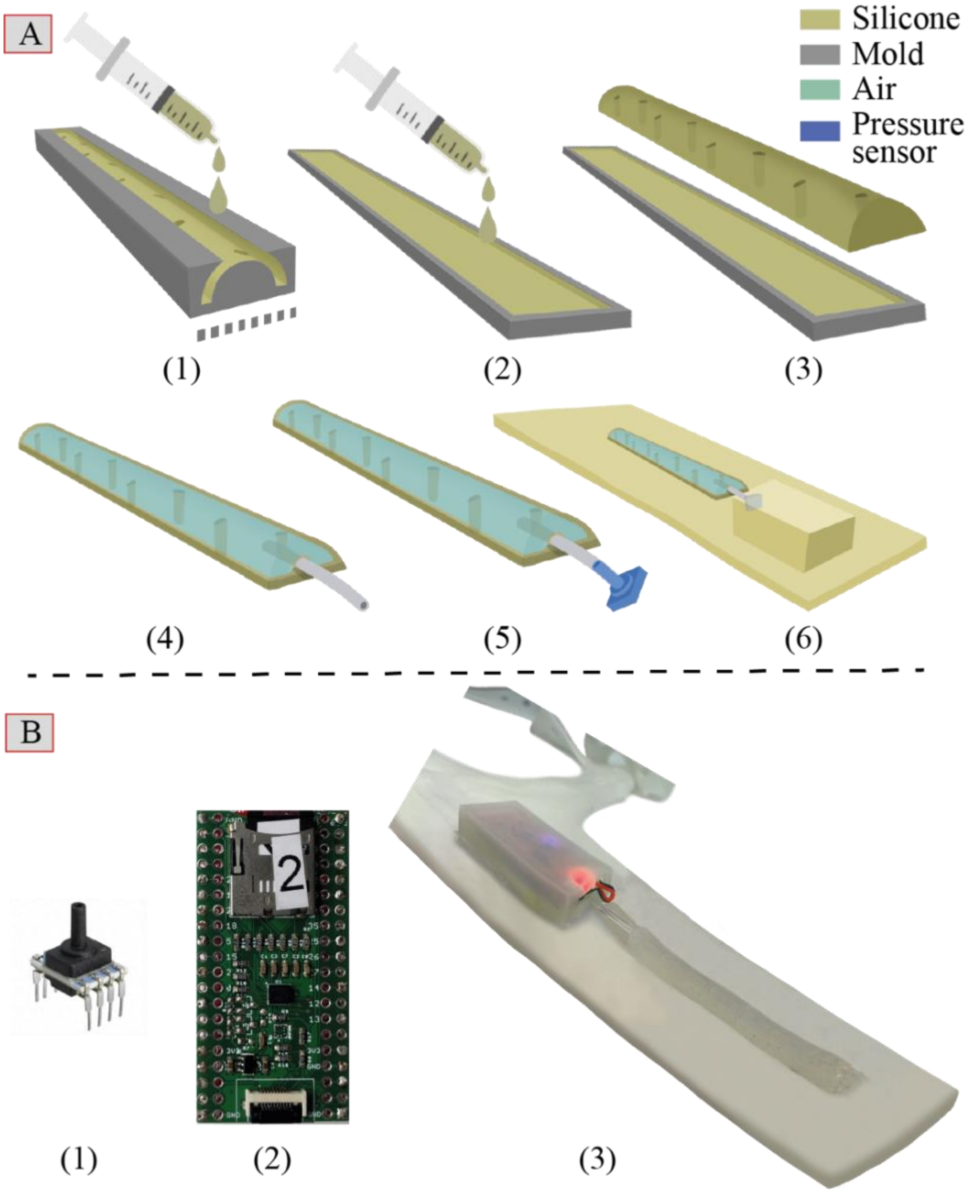


**Figure S1. A)** Illustrations of fabrication steps for the air-silicone composite transducer: **1.** Make the top part of the transducer by pouring liquid silicone into the mold and letting the silicone cure inside for 2 hours. **2.** Prepare the bottom part of the transducer by pouring liquid silicone into the mold that has a height of 1mm. **3.** Combine half-cured top part with the uncured bottom part. **4.** Place a thin silicone tube inside the air-silicone composite and adhere them using Sil-Poxy™ silicone adhesive. **5.** Place the gas pressure sensor inside the silicone tube. **6.** Attach gas pressure sensor to silicone harness. **B)** Images of the electronics and harness: **1.** Gas pressure sensor (HSCDLNN001PDAA3) **2.** PCB circuit for the collection, storing and transmission of the acquired data. **3.** Image of the complete system including the silicone harness, electronics, and air-silicone composite transducer.

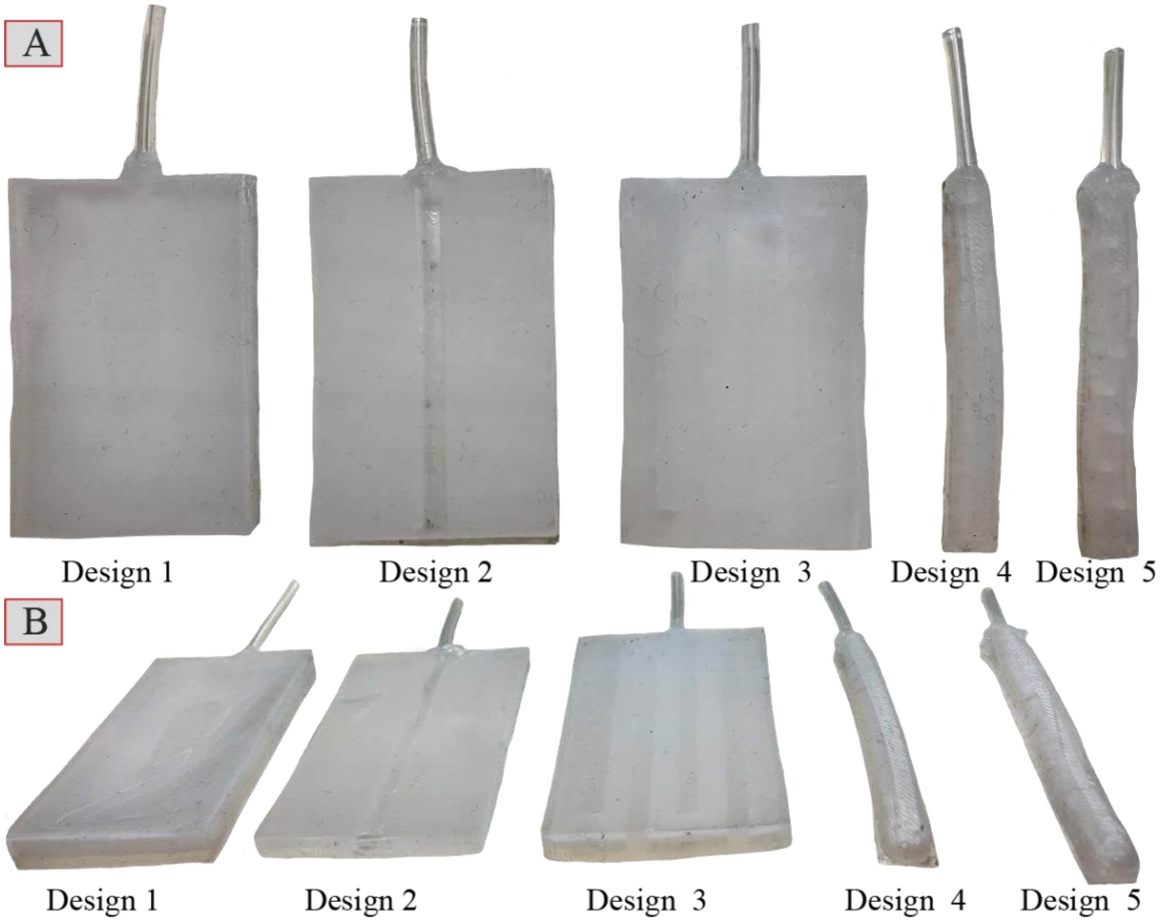


**Figure S2.** Image of ‘a’ variants of 5 main (Design 1 to Design 5) designs of the transducers. A) Top view. B)  Perspective view.

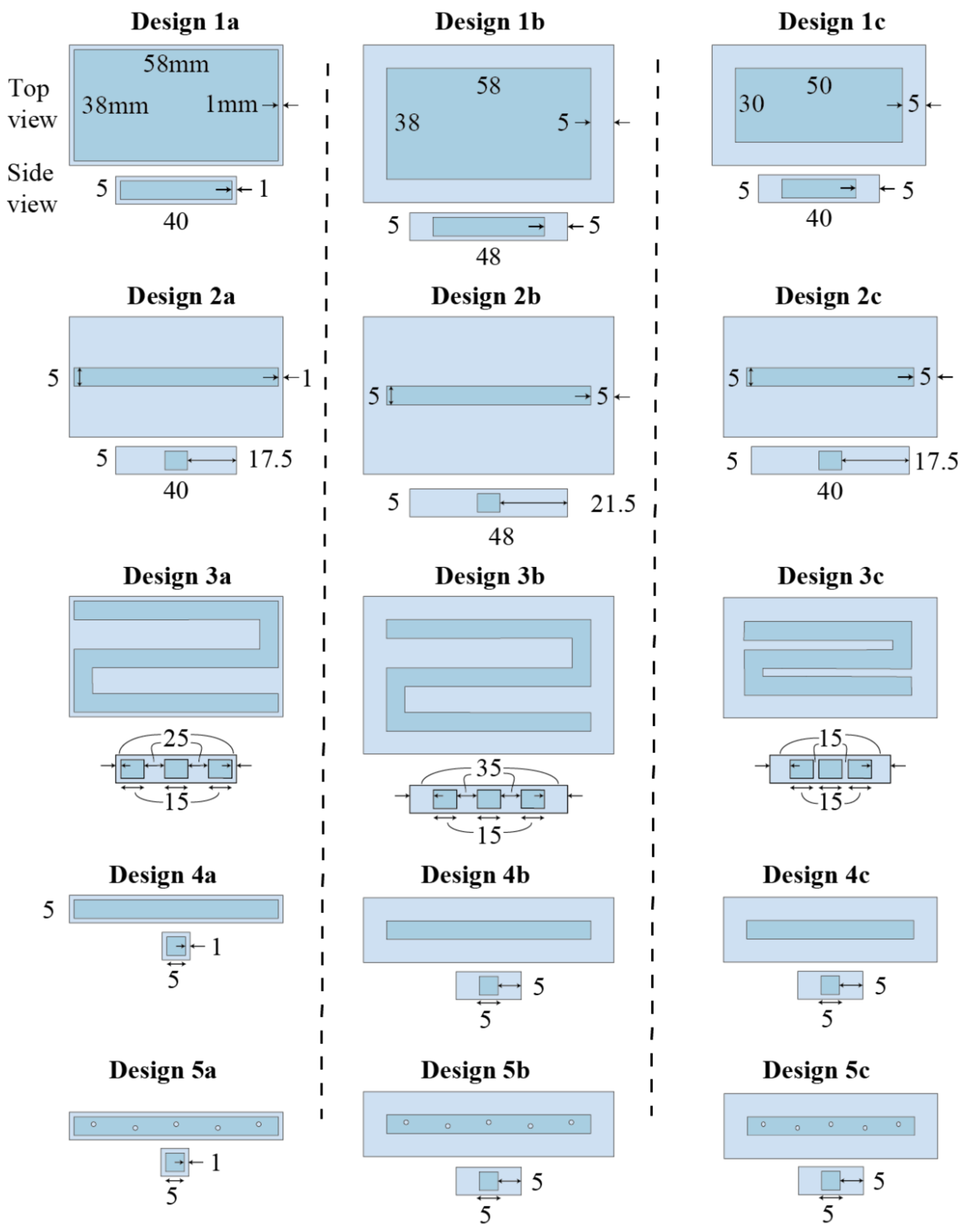


**Figure S3.** Top and side views of the 5 main designs of the transducers with 3 variants (a, b, and c) for testing. All transducers have the same height of 3mm. The first sample (Design 1) is composed of a large air tube (38mm width) encased with a thin layer (1mm) of silicone. The second sample (Design 2) is composed of a tiny air tube (5mm width) encased with a very thick layer (25mm) of silicone. The third sample (Design 3) is composed of three tiny air tubes (5mm width each) that are connected to each other and encased with a layer of silicone. The fourth sample (Design 4) is composed of a tiny air tube (5mm width) encased with a thin layer (1mm) of silicone. The fifth sample (Design 5) has the same shape as the fourth design, but it contains small silicone columns to reduce the amount of air (8% reduction) and to provide support against excessive pressure on the transducer. All designs explained above are originally fabricated with a silicone wall thickness of 1mm (a). Another variant (b) is created by increasing the wall thickness to 5mm. A final variant (c) is designed by decreasing the length of composite from 70mm to 60mm to reduce the air volume by more than 15%. The variants ‘b’ and ‘c’ were incorporated to explore the effect of silicone wall thickness and air volume on recording sensitivity, respectively. In addition, composites with increased heights of the air cavity of 3mm and 5mm were designed to find the height of the transducer that is most sensitive to chest motions.

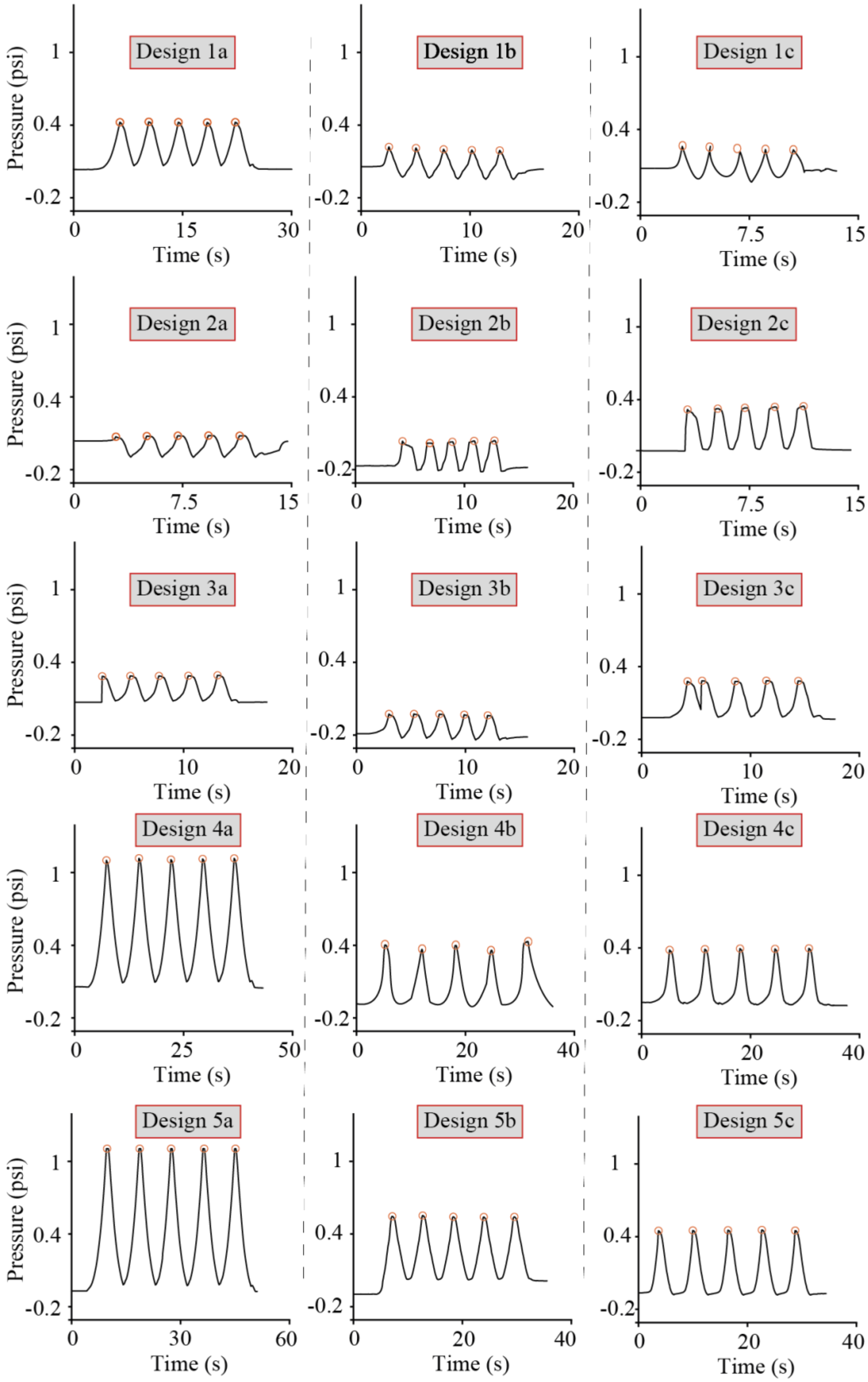


**Figure S4.** Recordings from the controlled experiments. The figure shows the response of our transducers (with a height of **3mm**) against an applied force ranging from 0 to 5N. Descriptions for designs can be found in Figure S3.

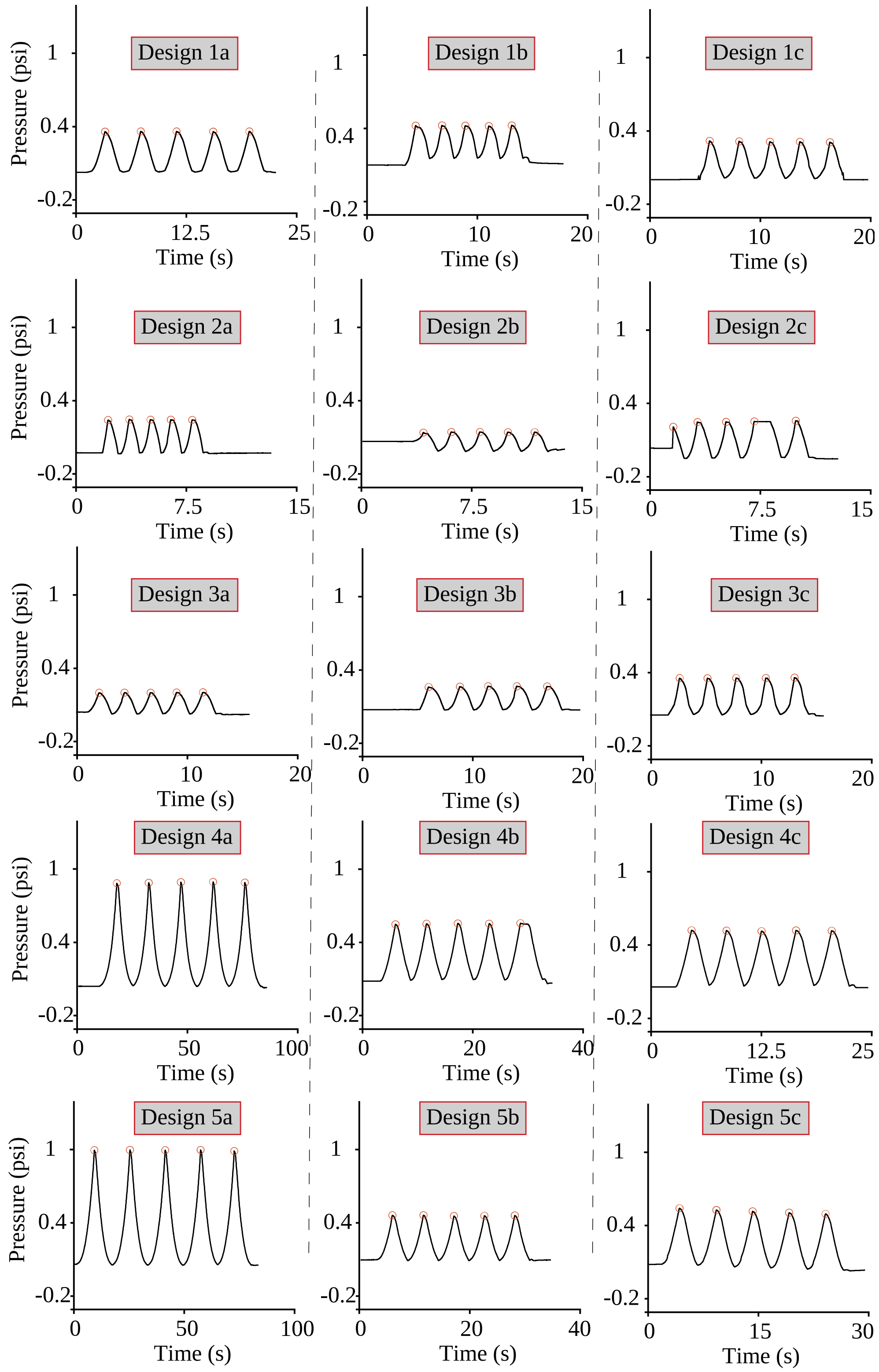


**Figure S5.** Recordings from the controlled experiments. The figure shows the response of our transducers (with a height of **5mm**) against an applied force ranging from 0 to 5N. Descriptions for designs can be found in Figure S3.

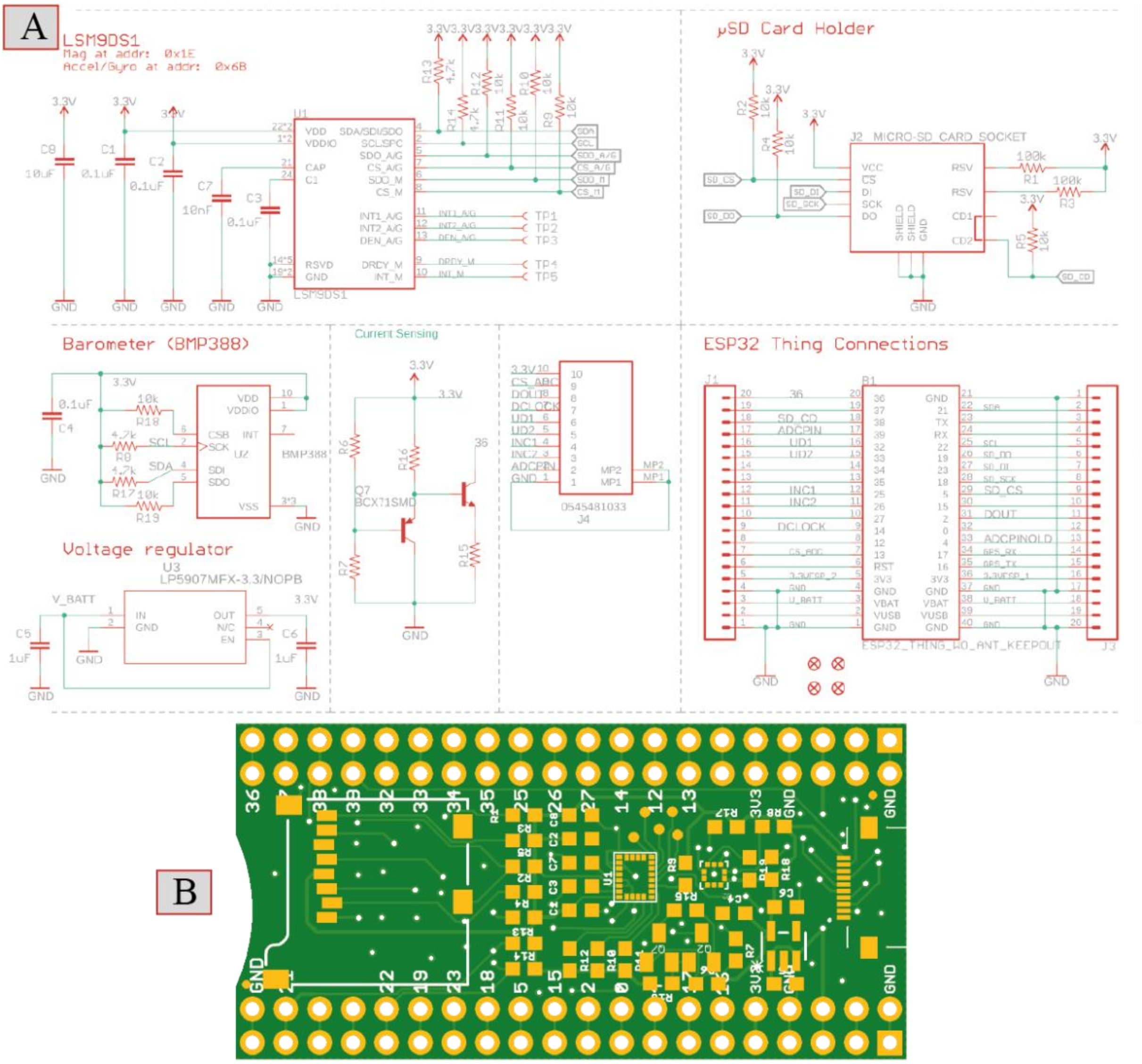


**Figure S6.** Electronics design for the collection, transmission and storage of the signals. A) Schematic design which includes a 3-axis accelerometer and barometer for monitoring of motion, SD card holder for storing data, voltage regulator for converting battery voltage to 3.3V, and pin heads for the connection to the ESP32 microcontroller used. B) Designed PCB for the system.

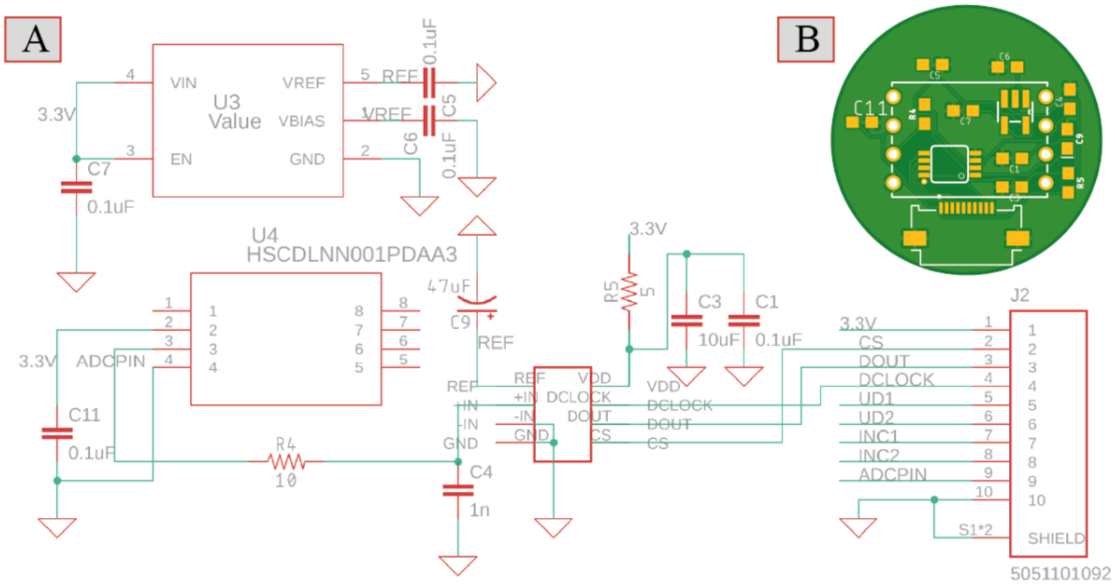


**Figure S7.** Data acquisition system for the gas pressure transducer. A) Schematic design of the data acquisition system, which includes high-precision voltage regulator, 16-bit ADC converter and the gas pressure sensor. B) Designed PCB for the system.

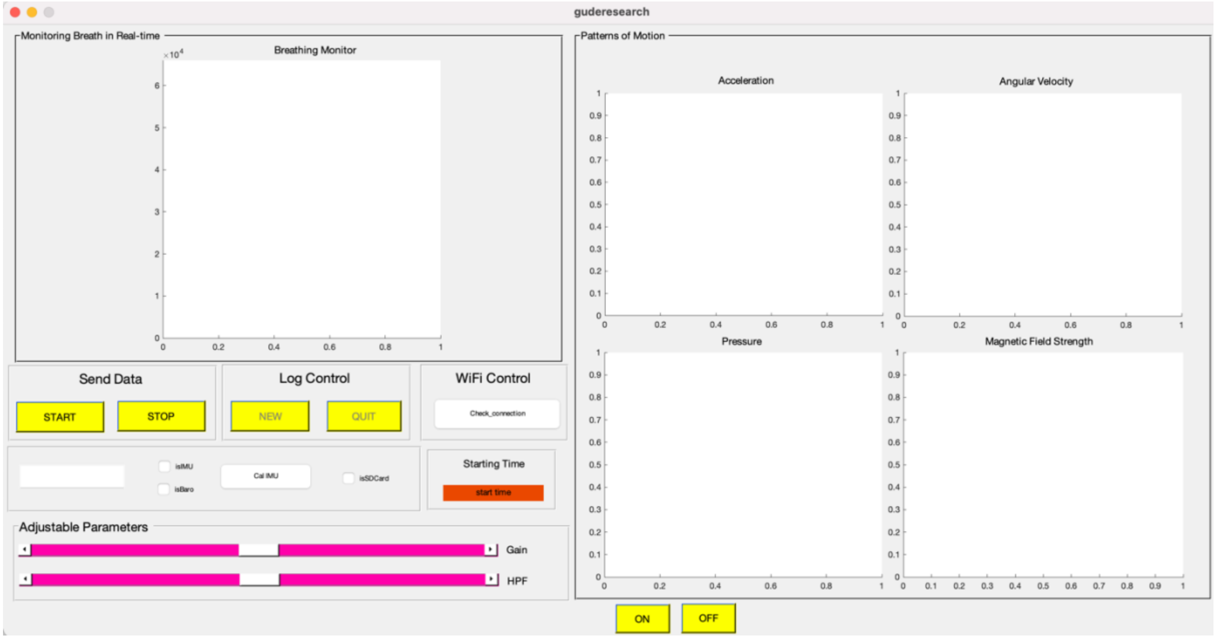


**Figure S8.** MATLAB interface to plot and process the acquired signals (breathing and motion) through Wi-Fi transmission. The MATLAB program is capable of remotely controlling the microcontroller by sending commands, including starting and stopping recordings, enabling, or disabling the acquisition of motion data, storing of collected data to the SD card, and labelling recordings.

### Detailed description of methods

#### Section S1: Design considerations of air silicone composite transducer

The chest of humans and dogs do not have a flat surface; therefore, flexible sensors, such as the one presented here, are required to adjust to the shape of the body for optimal functioning. In addition, using a longer transducer can achieve greater surface contact to provide an even better conformation to the body. For this reason, we increased the length of the Design 5a transducer from 60mm to 100mm (60 percent longer) to cover more surface on the chest wall. Increasing the length of the transducer, however, causes an increase in the air volume and decreases the sensitivity of the transducer, impeding the ability to catch small changes in the patterns of breathing such as rapid breath in humans or panting in dogs. For this purpose, to reduce the air volume of the transducer and restore sensitivity, we reduced the height of the composite to 2mm. Reduction of the height beyond 2mm will cause blockages in the transducer to chest motions and result in insensitive responses; thus, the volume of the transducer is further miniaturized by changing its shape to cylindrical instead of a cubic shape. As a result, the ASiT is able to capture tiny breathing movements due to its longer design that touches a greater area on the chest and smaller volume.

#### Section S2: Custom peak-detection algorithm for measuring breathing rate

In the peak detection, we utilized MATLAB's 'findpeaks' algorithm. From the MATLAB documentation: "findpeaks(data) returns a vector with the local maxima (peaks) of the input signal vector, data. A local peak is a data sample that is either larger than its two neighbouring samples or is equal to Inf. Non-Inf signal endpoints are excluded. If a peak is flat, the function returns only the point with the lowest index." (Signal Processing Toolbox: For use with MATLAB R2020b Update 2 (9.9.0.1524771). The MathWorks Inc.: Natick, Massachusetts, United States, 2020.)

This function only detects the high peaks; however, we set the parameters of the function to satisfy requirements for analysing of breathing signals such as the detection of low peaks (the point where exhalation ends and inspiration starts), and prediction of breathing rate in high accuracy. Moreover, we can determine the time spent for inhalation and exhalation. Besides detection of breathing rates, since the amplitude at low and high peaks are also known, we can extract more information from the breathing records (i.e., the speed of inhalation and exhalation). The sensitivity of the breathing detection algorithm can also be tuned depending on the peak distance (i.e., distance between successive peaks) and prominence (i.e., difference between peak and neighbouring points) parameters. Peak prominence is set to the 40% of the distance between the points of global maximum and global minimum. Peak distance is set to 0.2secs (the highest breathing rate can be detected is 1/0.2*60 = 300 breaths per minute).

#### Section S3: Extracting information from the patterns of breathing

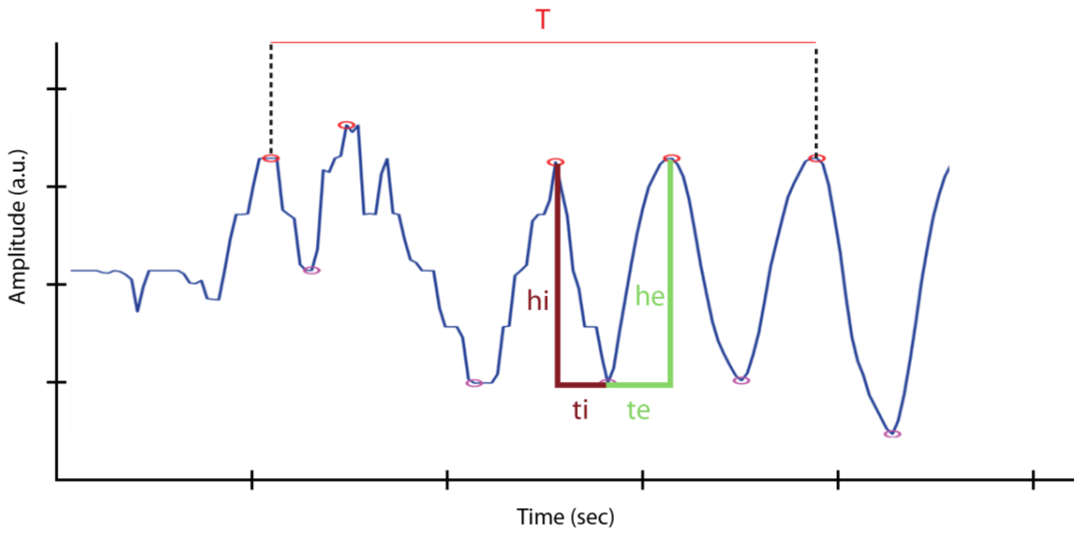


**Figure S9.** Parameters used to measure various patterns of breathing. 'T' represents the time between the first and the last high peaks. '$h_e$' represents the amplitude of exhalation (high peak(i) – low peak(i)) and '$h_i$' represents the amplitude of inhalation (high peak(i) – low peak(i+1)). '$t_e$' represents the time spent for one exhalation event and '$t_i$' represents the time spend for one inhalation event.

$$\text{Breaths per minute (br)} = \frac{\text{number of high peaks}}{\text{T}} \times 60$$

$$\text{Inhalation Speed (is)} = \frac{h_i}{t_i}$$

$$\text{Exhalation Speed (es)} = \frac{h_e}{t_e}$$

$$\text{Another breathing metric} = \frac{\text{Inhalation}}{\text{Exhalation}} = \frac{\text{is}}{\text{es}}$$

### Videos

**Video V1.** Real-time monitoring of breathing patterns of a detection dog during the training.

**Video V2.** Real-time monitoring of breathing patterns of a rat under anesthesia.
